# An *O*-acetylated derivative of piericidin A1 produced by *Kitasatospora* sp. A2-31 has potent activity against the cacao mirid bug, *Helopeltis bakeri* Poppius

**DOI:** 10.1101/2025.05.13.653897

**Authors:** Yousef Dashti, Felaine Anne Sumang, Emmanuel L.C. de los Santos, Alan Ward, Divina Amalin, Edwin P. Alcantara, Gregory L Challis

**Author notes:** School of Medical Sciences, Faculty of Medicine and Health, University of Sydney, Sydney NSW 2015, Australia (F.A.S.); UCB Pharma Ltd, 208 Bath Road, Slough, Berkshire, SL1 3WE, United Kingdom (E.L.C.d.l.S.).

## Abstract

There is an increasing demand for novel biopesticides to protect agricultural products and improve yields. To address this need, extracts from a library of Actinomycetes collected in the Philippines were evaluated against the cacao mirid bug, *Helopeltis bakeri* Poppius. Analysis of an active ethyl acetate extract from *Kitasatospora* sp. A2-31 led to the identification of a novel metabolite acetylpiericidin A1 (**3**) along with the known natural products piericidin A1 (**1**), piericidin A5 (**2**), chromomycin A2 (**4**), chromomycin A3 (**5**), and olivomycin A (**6**). Acetylpiericidin A1 demonstrated 100 % mortality against *H. bakeri*, while other metabolites exhibited either weak or no activity. Whole genome sequencing followed by bioinformatics analysis identified a gene downstream of the piericidin biosynthetic gene cluster that encodes a putative acetyltransferase proposed to catalyze acetylation of the C10 hydroxyl group of piericidin A1. The involvement of this gene in acetylpiericidin A1 biosynthesis was confirmed by (i) introducing an additional copy under the control of the *ermE\** promoter into *Kitasatospora* sp. A2-31, resulting in elevated production levels and (ii) through an *in vitro* enzymatic assay of the corresponding purified recombinant enzyme.

## Introduction

With the ever-expanding human population, crop production and food availability must increase significantly to meet global demand. Advances in agriculture, including improved pest management strategies, have contributed to enhanced crop yields.^1^ Currently, pest management heavily relies on the intensive use of synthetic pesticides.^2^ However, biopesticides have gained significant attention as a more sustainable alternative. Pesticidal specialized metabolites, categorized as biochemical biopesticides, possess several desirable attributes, including target specificity, low environmental persistence, and minimal toxicity to non-target organisms.^3, 4^ Actinomycetes have emerged as a promising source of natural product-based biopesticides. Several Actinomycete-derived compounds, such as spinosyns, blasticidin, bialaphos, abamectins, validamycin, milbemectin, streptomycin and fenpicoxamid, have already been commercialized as pesticides.^5^ Moreover, a recent surge in the availability of Actinomycete genome sequences, combined with advances in bioinformatics, has unveiled the untapped potential of this bacterial order to produce bioactive specialized metabolites, including novel biopesticides.^6-11^

The cacao mirid bug (*H. bakeri* Poppius, Hemiptera: Miridae; Figure S1), is a major insect pest of *Theobroma cacao* L. in Southeast Asia.^12, 13^ This bug not only damages cacao pods and young shoots by sucking sap for nutrition but can also serve as a carrier of pathogenic viruses during feeding.^14, 15^ With a worldwide distribution, cacao mirid bugs are reported to cause cacao yield losses ranging from 5 to 20%.^16^ Among the mirid bug pests associated with cacao, *Helopeltis* spp. are particularly well-recognized by farmers and agriculturalists in Southeast Asia as a serious threat. They cause characteristic lesions (Figure S1) and texture changes in the pods, which are readily observable symptoms of sap-feeding. In the Philippines, particularly in cacao-growing regions on the island of Luzon, the primary pest species is *H. bakeri*. Like other *Helopeltis* spp., it prefers feeding on young shoots (often causing shoot dieback under severe infestation) and on young or maturing pods. Lesions on young pods caused by sap-feeding can facilitate secondary infections by pathogenic microorganisms, leading to pod wilting.^13^ Here, we describe the discovery and biosynthesis of a novel specialized metabolite from *Kitasatospora* sp. A2-31 with potent activity against *H. bakeri*.

### Results and discussion

Crude extracts from a library of Actinomycetes isolated from various environments in the Philippines were tested for activity against the cacao mirid bug, *H. bakeri*. An ethyl acetate extract from a culture of *Kitasatospora* sp. A2-31, isolated from a gold mine tailing, exhibited potent activity. Profiling of the extract using UPLC-ESI-Q-TOF-MS revealed metabolites with molecular formulae corresponding to piericidin A1 (**1**),^17^ piericidin A5 (**2**),^18^ chromomycin A2 (**4**),^19, 20^ chromomycin A3 (**5**),^19, 20^ and olivomycin A (**6**),^20^ along with a novel compound. To identify the active component(s), and elucidate the structure of the new compound, semi-preparative HPLC was performed on an ethyl acetate extract from a two-liter culture of *Kitasatospora* sp. A2-31. The structures of the compounds with molecular formulae corresponding to known metabolites were confirmed by ^1^H and, where necessary, 2D NMR spectroscopy. The structure of the new compound **3** was elucidated using a combination of HR-MS and NMR spectroscopy (Figure 1, Figures S3-S7, Table S1). The molecular formula of C_27_H_40_NO_5_ deduced from HR-MS, suggested that this compound is a piericidin derivative. Detailed inspection of 2D NMR data (Figure S2 and Table S1) confirmed this hypothesis. The ^1^H NMR spectrum of compound **3** was very similar to that of piericidin A1 (**1**), except for an additional signal at 1.87 ppm, integrating for 3 protons. Similarly, the ^13^C NMR spectrum showed additional signals at 20.4 and 169.5 ppm, corresponding to methyl and carbonyl carbons, respectively. An HSQC correlation between the *δ*_H_ 1.87 ppm and *δ*_C_ 20.4 ppm signals, and an HMBC correlation between *δ*_H_ 1.87 ppm and *δ*_C_ 169.5 ppm signals indicated the compound is an acetylated derivative of piericidin A1. An HMBC correlation between the C10 proton and the carbonyl carbon of the acetyl group confirmed that the acetyl group is appended to the C10 hydroxyl group (Figure 1).

**Figure 1.**
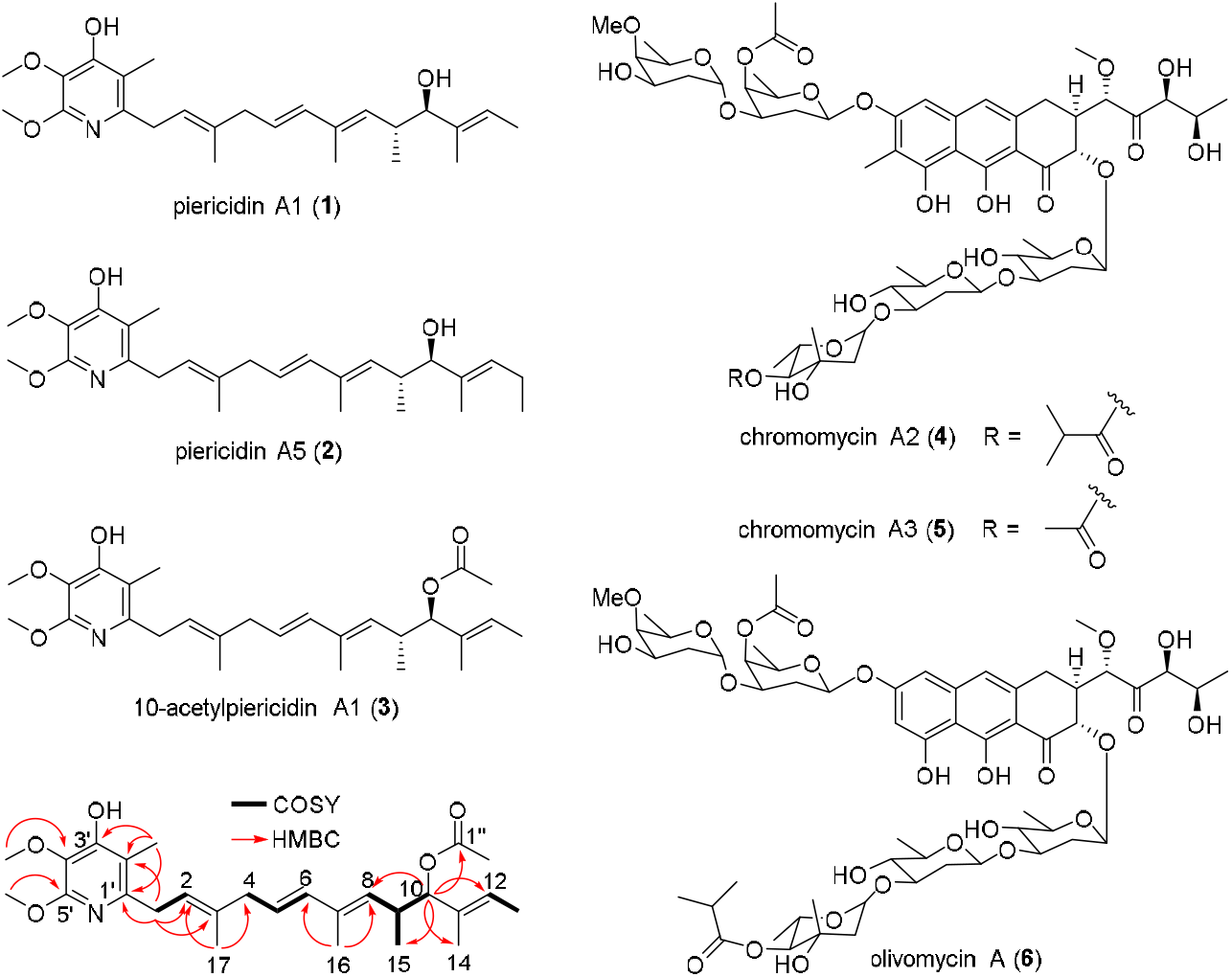
Structures of piericidin A1 (**1**), piericidin A5 (**2**), acetylpiericidin A1 (**3**), chromomycin A2 (**4**), chromomycin A3 (**5**), and olivomycin A (**6**) isolated from ethyl acetate extracts of *Kitasatospora* sp. A2-31. COSY and key HMBC correlations observed for the novel metabolite 10-acetylpiericidin A1 (**3**) are illustrated.

Purified **1**-**6** from *Kitasatospora* sp. A2-31 were screened for activity against *H. bakeri*. Acetylpiericidin A1 (**3**) caused 100% mortality, whereas only 10% mortality was observed with piericidin A1 (**1**), and the other compounds were inactive. The estimated median lethal dose (LD_50_) of acetylpiericidin A1 (**3**) to third-instar *H. bakeri* was 335 ng/insect. Piericidin analogues are produced by various *Streptomyces* strains.^17, 18, 21-23^ Piericidin A1 (**1**) has been reported to possess antibacterial and antifungal activity, in addition to selective insecticidal activity,^24, 25^ and is also a selective antitumor agent.^26^ The antimicrobial and insecticidal activity of piericidin A1 (**1**) are attributed to its structural similarity to ubiquinone (coenzyme Q10). It is a potent competitive inhibitor of both bacterial and mitochondrial NADH-ubiquinone oxidoreductase (complex I).^24, 25^ Earlier studies showed that the hydroxyl group attached to the pyridine, which mimics the quinone structure of ubiquinone, is essential for the activity of piericidins.^27^ Structure-activity relationship investigations employing piericidin analogues revealed that variations in the alkyl side chain structure do not significantly impact activity, because derivatives with even simple alkyl chains retain potent activity.^28^ It has been suggested that the methyl-branched tetraene side chain enhances the hydrophobicity of the piericidins.^29^ The strong insecticidal activity of acetylpiericidin A1 (**3**) compared to piericidin A1 (**1**) may simply be a result of increased hydrophobicity due to C10-acetylation, which is likely to enhance cell wall permeability and target accessibility in *H. bakeri* relative to piericidin A1 (**1**). Further studies will be needed to establish the molecular basis for the dramatic difference in potency between **1** and **3**.

To identify the putative acetylpiericidin A1 biosynthetic gene cluster (BGC), we sequenced and assembled the complete genome of *Kitasatospora* sp. A2-31 using a combination of Illumina and Oxford Nanopore technologies. Bioinformatics analyses identified the *api* cluster of genes (NCBI accession number: PP459019), which exhibits a high degree of similarity to the piericidin A1 (**1**) BGCs found in *Streptomyces piomogeues* var. Hangzhouwanensis and *Streptomyces* sp. SCSIO 03032 (Table S2).^30, 31^ Eleven genes in the *api* BGC are proposed to encode piericidin A1 (**1**) biosynthetic enzymes, including six modular polyketide synthase (PKS) subunits (ApiA1-ApiA6), an amidotransferase (ApiD), a putative cyclase (ApiC), a monooxygenase (ApiE), and two methyltransferases (ApiB1 and ApiB2) (Figure 2 and Table S2). A twelfth gene, *apiR*, encodes a SARP-family transcriptional regulator (Figure 2 and Table S2).^30-32^

**Figure 2.**
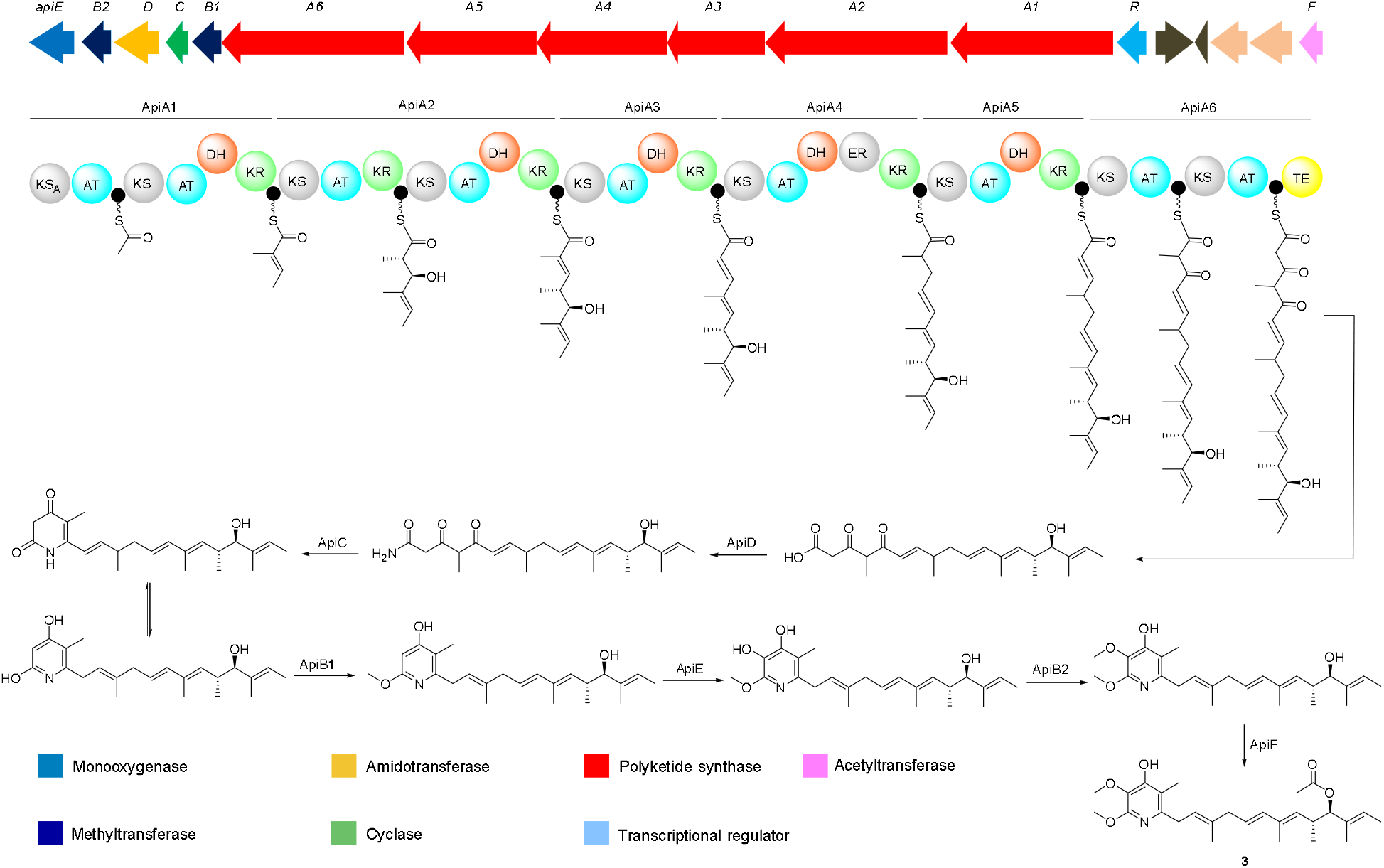
Organization of the acetylpiericidin A1 (**3**) BGC in *Kitasatospora* sp. A2-31 and proposed pathway for acetylpiericidin A1 (**3**) assembly.

The modular PKS subunits (ApiA1-ApiA6) are proposed to assemble the carbon skeleton of piericidin A1 (**1**) from four malonyl-CoA and five methyl malonyl-CoA units. The fully assembled polyketide chain is released from the assembly line by hydrolysis catalyzed by the thioesterase (TE) domain appended to the C-terminus of the final PKS module. The putative ATP-dependent amidotransferase (ApiD) is proposed to catalyze conversion of the PKS product to the corresponding amide, and a putative cyclase (ApiC) is hypothesized to accelerate the spontaneous cyclisation of the amide to an α-pyridone.^30^ Two *O*-methyltransferases (ApiB1 and ApiB2) and a FAD-dependent monooxygenase (ApiE) are proposed to convert the α-pyridone intermediate to piericidin A1 (**1**). Consistent with this hypothesis, the ApiE homologue PieE has been shown to hydroxylate the 4□-position of the α-pyridone in *Streptomyces* sp. SCSIO 03032 and PieB2, a homologue of ApiB2, catalyzes methylation of the resulting hydroxyl group.^31^

In addition to homologs of piericidin A1 (**1**) biosynthetic genes in *Streptomyces piomogeues* var. Hangzhouwanensis and *Streptomyces* sp. SCSIO 03032, *apiF* was identified in the right flank of the *api* BGC in *Kitasatospora* sp. A2-31 (Figure 2). We hypothesized this gene encodes a putative acyltransferase (31% sequence identity to spermidine *N*-acetyltransferase SpeG) that catalyzes acetylation of the piericidin A1 C10 hydroxyl group, affording acetylpiericidin A1 (**3**). To investigate this hypothesis, ApiF was overproduced in *E. coli* as an N-terminal His_6_-fusion protein and purified to homogeneity using nickel affinity chromatography. The integrity and identity of the purified protein was confirmed by intact protein mass spectrometry (Figure S8). Overnight incubation of purified recombinant ApiF with piericidin A1 and acetyl coenzyme A at room temperature resulted in the formation of acetylpiericidin A1 (**3**) (Figure 3). Moreover, introduction of a second copy of *apiF* into *Kitasatospora* sp. A2-31, under the control of the constitutive *ermE*\* promoter, doubled the yield of acetylpiericidin A1 (**3**) (from 0.2 to 0.4 mg/L).

**Figure 3.**
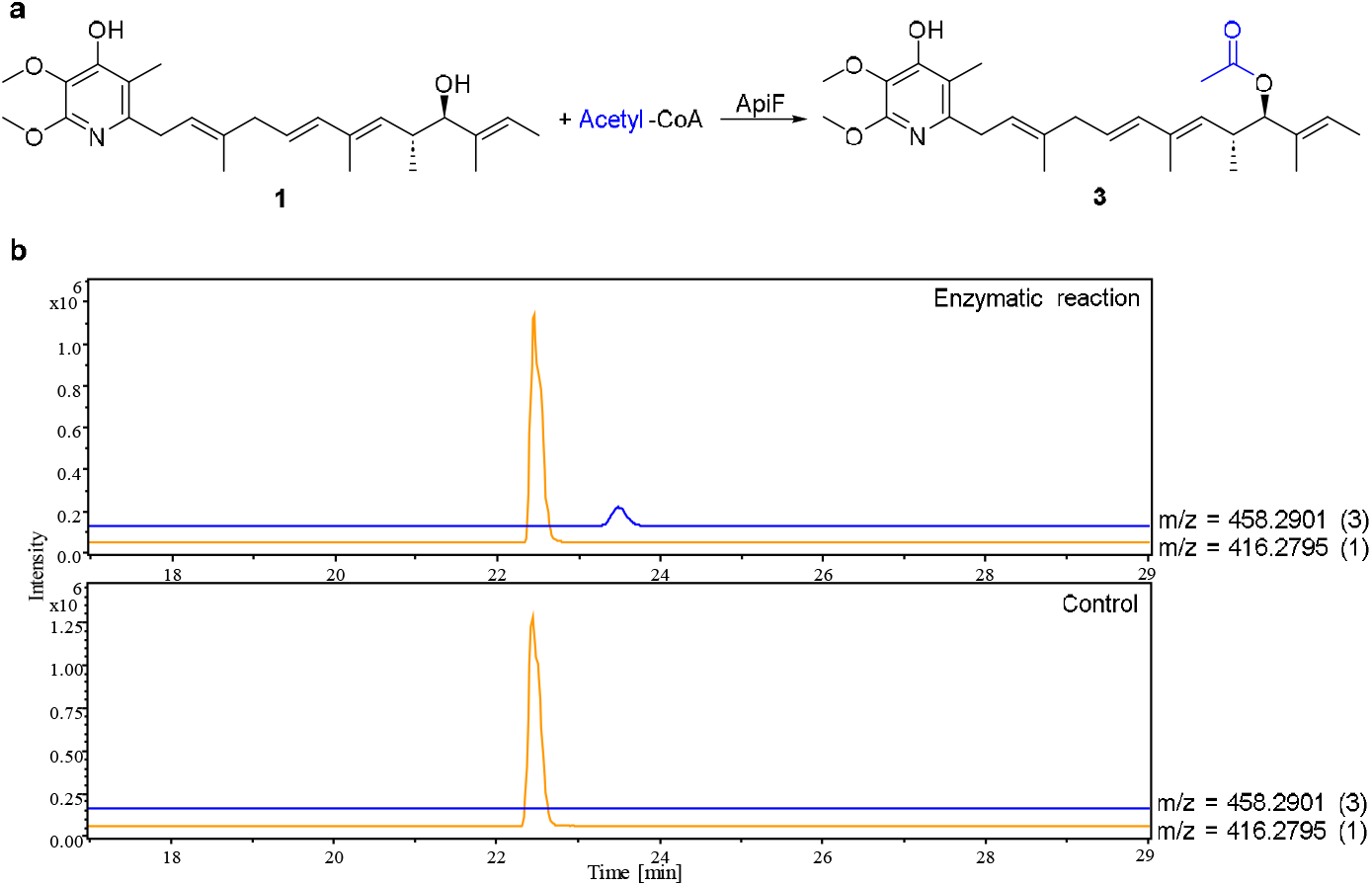
ApiF catalyzes the conversion of piericidin A1 (**1**) to acetylpiericidin A1 (**3**). **a**. Reaction catalyzed by ApiF. **b**. Extracted ion chromatograms at *m/z* = 416.2795 and 458.2901, corresponding to [M+H]^+^ for **1** and **3**, respectively, from UHPLC-ESI-Q-TOF MS analyses of enzymatic and negative control reactions.

In conclusion, screening of culture extracts of Actinomycetes isolated from diverse environments in the Philippines led to the discovery of acetylpiericidin A1 (**3**), a novel specialized metabolite with potent activity against the cacao mirid bug, *H. bakeri*. The comparatively weak activity of piericidin A1 (**1**) suggests functionalization of the C10 hydroxyl group may be a promising strategy for enhancing the efficacy of this metabolite class against the cacao mirid bug. Identification of the acetylpiericidin A1 biosynthetic gene cluster in *Kitasatospora* sp. A2-31 enabled us to identify *apiF* as the gene encoding the acetyl transferase responsible for converting piericidin A1 (**1**) to acetyl piericidin A1 (**3**). Although ApiF appears to have only modest catalytic activity, directed evolution could be used to improve this and broaden its substrate scope, enabling it to be developed into a useful biocatalyst for production of various acetylpiericidin A1 analogues. While acetylation of the C10 hydroxyl in piericidin A1 (**1**) may simply enhance cell wall permeability and therefore target accessibility, further experiments will be required to illuminate the molecular basis for the dramatic enhancement in activity of acetylpiericidin A1 against *H. bakeri*.

## Materials and Methods

### General experimental procedures

UHPLC-ESI-Q-TOF-MS analyses were performed on a Dionex UltiMate 3000 UHPLC connected to a Zorbax Eclipse Plus C18 column (100 × 2.1 mm, 1.8 μm) coupled to a Bruker MaXis Impact mass spectrometer. Both mobile phases, water (A) and acetonitrile (B), were supplemented with 0.1% formic acid. The LC method employed a gradient run from 5 to100% B over 30 minutes at a flow rate of 0.2 mL/min. The mass spectrometer was operated in positive ion mode with a scan range of 50-3000 *m/z*. A solution of 1 mM sodium formate was used for calibration through a loop injection of 20 μL at the start of the run. NMR spectra were recorded on a Bruker Avance III 600 MHz spectrometer in DMSO-*d*_*6*_. ^1^H and ^13^C NMR chemical shifts were referenced to the residual protiated solvent peaks at *δ*_H_ 2.50 and *δ*_C_ 39.51.

### Strain isolation

Fifteen mine tailings were collected from different locations within a private gold mining corporation located at Baguio City, Philippines. For each mine tailing, one gram of soil was used to isolate Actinomycetes by performing four tenfold serial dilutions, mixing using a vortex mixer. Subsequently, 100 μl from both 10^−2^ and 10^−3^ dilutions were plated on separate YEME agar plates (10 g/L glucose, 5 g/L peptone, 3 g/L malt extract, 3 g/L yeast extract) and the resulting cultures were incubated at 30 □ for three days. Potential Actinomycete colonies were picked and streaked onto fresh YEME agar plates to obtain pure single colonies.

### Production, extraction and HPLC purification of metabolites from *Kitasatospora* sp. A2-31

*Kitasatospora sp*. A2-31 was grown on 2 L of ISP2 agar medium (4 g/L glucose, 4 g/L yeast extract, 10 g/L malt extract, 2 g/L CaCO_3_, 15g/L Bacto agar) for six days at 30 °C. The agar was extracted twice with EtOAc and the combined extracts were evaporated to dryness. The residue was adsorbed onto dental cotton, which was packed into a stainless steel HPLC guard cartridge (10 × 30 mm) connected to a semi-preparative C18 Betasil column (21.2 mm × 150 mm). HPLC separation was performed at a flow rate of 9 mL/min using the following elution profile: isocratic 5% acetonitrile for 5 min, followed by a linear gradient from 5 to 100% acetonitrile over 45 min, then isocratic 100% acetonitrile for 10 min. 120 fractions were collected at 30 second intervals over the 60 min run time. Chromomycin A3 eluted in fraction 67, olivomycin A in fraction 70, and chromomycin A2 in fractions 76 and 77. Pure piericidin A1, piericidin A5, and acetylpiericidin A1 were obtained in fractions 102, 107, and 108, respectively. The structures of known compounds were confirmed by comparisons with reported HR-MS and NMR data.

*Acetylpiericidin A1*: pale yellow oil (0.2 mg/L); ^1^H NMR (600 MHz, DMSO-*d*_*6*_) and ^13^C NMR (150 MHz, DMSO-*d*_*6*_), see Table S1; HRESIMS *m*/*z* 458.2905 [M + H]^+^ (calcd for C_27_H_40_NO_5_, 458.2901).

### Genomic DNA extraction

Spores of *Kitasatospora* sp. A2-31 were inoculated in 10 mL Tryptic Soy Broth (Pronadisa) and the resulting cultures we incubated at 30 □ with shaking at 200 rpm for three days. Genomic DNA was extracted from separated biomass using the Gentra® Puregene® Kit (Qiagen, Hilden, Germany) following the manufacturer’s protocol with some modifications. Instead of using 500 μl of cell culture, 100 mg of cell biomass was used for the extraction. The cells were macerated after addition of Cell Lysis Solution and incubation at 80 □ for five minutes. The isolated genomic DNA was resuspended in 50 μl hydration solution.

### Genome sequencing, assembly, error correction, and annotation

The genomic DNA of *Kitasatospora* sp. A2-31 was sequenced by the MicrobesNG DNA Sequencing Facility at the University of Birmingham using a combination of Illumina and Oxford Nanopore sequencing technologies. Illumina raw reads were trimmed using trim galore^33^ and assembled with SPAdes,^34^ then optimized using Unicycler.^35^ Nanopore raw reads were assembled separately with Flye^36^ and a hybrid assembly was performed with Unicycler.^35^ The resulting genome contigs were polished by mapping the Illumina reads back to the assembled contigs using Mira 5.0^33^ and Pilon.^37^ The putative acetylpiericidin A1 BGC was identified using antiSMASH.^38^ The identified BGC was further validated for coverage and identity by mapping both Illumina and nanopore reads and re-annotated using the NCBI Prokaryotic Genome Annotation Pipeline (PGAP).^39^ The sequence of the annotated BGC has been deposited in the NCBI (accession number: PP459019).

### Overproduction and purification of ApiF

*apiF* was amplified from genomic DNA of *Kitasatospora sp*. A2-31 using Q5 DNA polymerase (NEB) with primers ApiF-F (GACCATATGATGCTACAAGGCGCCCAC) and ApiF-R (GACAAGCTTTCAGCCCCGCCACTCCTC). PCR products were separated on a 1% agarose gel, and bands of the expected size were excised and purified using the GeneJET Gel Extraction Kit (Thermo Scientific). The purified PCR product was digested with *Nde*I and *Hin*dIII and subsequently ligated into the *Nde*I*/Hin*dIII-digested pET28a vector. The ligation mixture was used to transform *E. coli* TOP10 cells (Invitrogen), which were plated on LB agar containing kanamycin (50 μg/mL). To verify the integrity of the insert, selected colonies were cultured overnight in LB medium, plasmids were isolated using the GeneJET Plasmid Miniprep Kit (Thermo Scientific), and the inserts were sequenced using universal forward and reverse primers. A verified clone was used to transform *E. coli* BL21(DE3), and recombinant His6-ApiF was overproduced and purified as reported previously.^40^

### ApiF enzymatic assay for conversion of piericidin A1 to acetylpiericidin A1

A 200 μL reaction mixture containing piericidin A1 (100 μM), acetyl-CoA (100 μM), and ApiF (10 μM) in 25 mM Tris-HCl (pH 8.0) was incubated overnight at room temperature. The reaction was terminated by adding 200 μL of methanol. After centrifugation to remove precipitates, the supernatant was analyzed by UHPLC-ESI-Q-TOF-MS. For the negative control, ApiF was denatured by boiling at 100 °C for 15 min prior to incubation.

### Expression of *apiF* in *Kitasatospora* sp. A2-31

*apiF* was amplified from *Kitasatospora sp*. A2-31 genomic DNA using Q5 DNA polymerase (NEB) with the primers apiF-For (GACCATATGATGCTACAAGGCGCCCAC) and apiF-Rev (GACGAATTCTCAGCCCCGCCACTCCTC). The resulting amplimer was processed as described above and cloned into the *Nde*I*/Eco*RI sites of pIB139.^41^ In this vector, the *apiF* gene is placed under the control of the constitutive *ermE*\* promoter. After confirming the integrity of the insert by sequencing, the plasmid was used to transformed *E. coli* ET12567/pUZ8002 by electroporation and introduced into *Kitasatospora sp*. A2-31 via conjugation, as described.^42^ The engineered strain and wild type *Kitasatospora sp*. A2-31 were cultured separately on 1 L of ISP2 agar medium, and HPLC purification was used to compare acetylpiericidin A1 yields.

### Source of test insects

The *H. bakeri* used in this study were first- and second-generation insects from a stock population maintained in the insect rearing laboratory of the Biological Control Research Unit at De La Salle University - Science and Technology Complex, Biñan, Laguna, Philippines. The stock population was established using nymphs and adult mirid bugs were collected from infested cacao trees in Candelaria, Quezon (13.922866°N, 121.454605°E), Pili, Camarines Sur (13.583676°N, 123.264191°E), and Naga City, Camarines Sur (13.664177°N, 123.260979°E). Sweet potato (*Ipomea batatas* (L.)) was used as a feeding and oviposition medium for rearing *H. bakeri* in the laboratory.^43^ The shoots were found to be a suitable feeding medium for both nymphal instars and adult mirid bugs. Laboratory conditions were maintained at 26-28 °C and 59-73% relative humidity under a 12h on/12 h off photoperiod.

### Bioactivity Assays

Ethyl acetate and methanol extracts of Actinomycetes, resuspended in DMSO, were subjected to a brine shrimp lethality assay for preliminary toxicity screening.^44^ Brine shrimps were hatched by incubating approximately 1 g of eggs in 1 L of water containing 25–30 g of non-iodized salt, under continuous aeration and constant light. After 12 hours, the hatched nauplii were collected by light attraction and concentrated into a small vial. Ten nauplii were transferred into each well of a 96-well plate, with a final volume of 148 µl per well. Subsequently, 2 µl of the extract was added to each well, with DMSO serving as the negative control. Plates were incubated at room temperature under constant light. Toxicity was assessed after 12 hours by calculating the percentage of mortality. Extracts inducing 90–100% lethality were prioritized for further testing against *H. bakeri*.

A topical application method was used to assess insecticidal efficacy of purified metabolites against *H. bakeri*. Stock solutions of the metabolites were prepared in a buffer containing 0.05% DMSO and 0.1% Triton X-100. From these stock solutions, nine test concentrations ranging from 20 to 100 ng/µl were prepared. Buffer containing 0.05% DMSO and 0.1% Triton X-100 without metabolite was used as the negative control. For each test, 5 μL of the respective solution was applied to the dorsal side of the thoracic segment of each nymph. Ten third nymphal stage insects were used for each concentration. Mortality was scored after 24 hours, and the median lethal dose (LD_50_) was estimated using Priprobit software.^45^

## Author Contributions

G.L.C., E.L.C.d.l.S., Y.D. and E.P.A. designed and coordinated the project. E.L.C.d.l.S. and A.W. performed the genome assembly and bioinformatics analysis. Y.D. isolated and characterized the known and novel metabolites from *Kitasatospora* sp. A2-31, overproduced, purified and characterized ApiF, and expressed *apiF* in *Kitasatospora* sp. A2-31. F.A.S. and E.P.A. isolated *Kitasatospora* sp. A2-31 and grew cultures for initial biological testing. F.A.S. and D.A. tested the activity of culture extracts and purified compounds against *H. bakeri*. Y.D. and G.L.C. wrote the manuscript with input from the other authors.

## Notes

The authors declare the following competing financial interest(s): Gregory L. Challis is a co-founder, non-executive director, and shareholder of Erebagen Ltd. The other authors declare no competing financial interest.

## Acknowledgements

This project was funded by the British Council through a Newton Institutional Links Award (grant ref. 261846416). The Bruker MaXis Impact UHPLC-ESI-Q-TOF-MS instrument used in this research was purchased with a grant from the BBSRC (BB/K002341/1 to G.L.C.). E.L.C.d.l.S. was a Research Career Development Fellow in the Warwick Integrative Synthetic Biology Centre, which is supported by a grant from the BBSRC and EPSRC (BB/M017982/1). G.L.C. was the recipient of a Wolfson Research Merit Award from the Royal Society (grant no. WM130033).

## References

(1) Pretty, J. Agricultural sustainability: concepts, principles and evidence. Philos. Trans. R. Soc. B 2008, 363 (1491), 447–465. DOI: doi:10.1098/rstb.2007.2163.

(2) Chandler, D.; Bailey, A. S.; Tatchell, G. M.; Davidson, G.; Greaves, J.; Grant, W. P. The development, regulation and use of biopesticides for integrated pest management. Philos. Trans. R. Soc. B 2011, 366 (1573), 1987–1998. DOI: 10.1098/rstb.2010.0390.

(3) Hajek, A. E. Natural Enemies: An Introduction to Biological Control; Cambridge University Press, 2004. DOI: 10.1017/CBO9780511811838.

(4) Seiber, J. N.; Coats, J.; Duke, S. O.; Gross, A. D. Biopesticides: state of the art and future opportunities. J. Agric. Food Chem. 2014, 62 (48), 11613–11619. DOI: 10.1021/jf504252n.

(5) Copping, L. G.; Duke, S. O. Natural products that have been used commercially as crop protection agents. Pest. Manag. Sci. 2007, 63 (6), 524–554. DOI: 10.1002/ps.1378.

(6) Wilkinson, B.; Micklefield, J. Mining and engineering natural-product biosynthetic pathways. Nat. Chem. Biol. 2007, 3 (7), 379–386. DOI: 10.1038/nchembio.2007.7.

(7) Omura, S.; Ikeda, H.; Ishikawa, J.; Hanamoto, A.; Takahashi, C.; Shinose, M.; Takahashi, Y.; Horikawa, H.; Nakazawa, H.; Osonoe, T.; Kikuchi, H.; Shiba, T.; Sakaki, Y.; Hattori, M. Genome sequence of an industrial microorganism Streptomyces avermitilis: deducing the ability of producing secondary metabolites. PNAS 2001, 98 (21), 12215–12220. DOI: 10.1073/pnas.211433198.

(8) Gross, H. Strategies to unravel the function of orphan biosynthesis pathways: recent examples and future prospects. Appl. Microbiol. Biotechnol. 2007, 75 (2), 267–277. DOI: 10.1007/s00253-007-0900-5.

(9) Bentley, S. D.; Chater, K. F.; Cerdeño-Tárraga, A. M.; Challis, G. L.; Thomson, N. R.; James, K. D.; Harris, D. E.; Quail, M. A.; Kieser, H.; Harper, D.; Bateman, A.; Brown, S.; Chandra, G.; Chen, C. W.; Collins, M.; Cronin, A.; Fraser, A.; Goble, A.; Hidalgo, J.; Hornsby, T.; Howarth, S.; Huang, C. H.; Kieser, T.; Larke, L.; Murphy, L.; Oliver, K.; O’Neil, S.; Rabbinowitsch, E.; Rajandream, M. A.; Rutherford, K.; Rutter, S.; Seeger, K.; Saunders, D.; Sharp, S.; Squares, R.; Squares, S.; Taylor, K.; Warren, T.; Wietzorrek, A.; Woodward, J.; Barrell, B. G.; Parkhill, J.; Hopwood, D. A. Complete genome sequence of the model actinomycete Streptomyces coelicolor A3(2). Nature 2002, 417 (6885), 141–147. DOI: 10.1038/417141a.

(10) Challis, G. L. Genome mining for novel natural product discovery. J. Med. Chem. 2008, 51 (9), 2618–2628. DOI: 10.1021/jm700948z.

(11) Wilson, M. C.; Mori, T.; Ruckert, C.; Uria, A. R.; Helf, M. J.; Takada, K.; Gernert, C.; Steffens, U. A.; Heycke, N.; Schmitt, S.; Rinke, C.; Helfrich, E. J.; Brachmann, A. O.; Gurgui, C.; Wakimoto, T.; Kracht, M.; Crusemann, M.; Hentschel, U.; Abe, I.; Matsunaga, S.; Kalinowski, J.; Takeyama, H.; Piel, J. An environmental bacterial taxon with a large and distinct metabolic repertoire. Nature 2014, 506 (7486), 58–62. DOI: 10.1038/nature12959.

(12) Stonedahl, G. M. The Oriental species of Helopeltis (Heteroptera: Miridae): a review of economic literature and guide to identification. Bull. Entom. Res. 1991, 81 (4), 465–490. DOI: 10.1017/S0007485300032041.

(13) Amalin, D. M.; Averion, L.; Bihis, D.; Legaspi, J. C.; David, E. F. Effectiveness of Kaolin Clay Particle Film in Managing Helopeltis collaris (Hemiptera: Miridae), a Major Pest of Cacao in the Philippines. Fla. Entomol. 2015, 98 (1), 354–355, 352. DOI: 10.1653/024.098.0156.

(14) Muhamad, R.; Way, M. J. Damage and crop loss relationships of Helopeltis theivora, Hemiptera, Miridae and cocoa in Malaysia. Crop Prot. 1995, 14 (2), 117–121. DOI: 10.1016/0261-2194(95)92865-K.

(15) Rice, R. A.; Greenberg, R. Cacao cultivation and the conservation of biological diversity. Ambio 2000, 29 (3), 167–173. DOI: 10.1579/0044-7447-29.3.167.

(16) Gotsch, N. Cocoa crop protection: an expert forecast on future progress, research priorities and policy with the help of the Delphi survey. Crop Prot. 1997, 16 (3), 227–233. DOI: 10.1016/S0261-2194(96)00099-3.

(17) Takahashi, N.; Suzuki, A.; Tamura, S. Structure of Piericidin A. J. Am. Chem. Soc. 1965, 87 (9), 2066–2068. DOI: 10.1021/ja01087a050.

(18) Kroiss, J.; Kaltenpoth, M.; Schneider, B.; Schwinger, M.-G.; Hertweck, C.; Maddula, R. K.; Strohm, E.; Svatoš, A. Symbiotic streptomycetes provide antibiotic combination prophylaxis for wasp offspring. Nat. Chem. Biol. 2010, 6, 261. DOI: 10.1038/nchembio.331.

(19) Miyamoto, M.; Kawamatsu, Y.; Kawashima, K.; Shinohara, M.; Tanaka, K.; Tatsuoka, S.; Nakanishi, K. Chromomycin A2, A3 and A4. Tetrahedron 1967, 23 (1), 421–437. DOI: 10.1016/S0040-4020(01)83328-7.

(20) Yoshimura, Y.; Koenuma, M.; Matsumoto, K.; Tori, K.; Terui, Y. NMR studies of chromomycins, olivomycins, and their derivatives. J. Antibiot. 1988, 41 (1), 53–67. DOI: 10.7164/antibiotics.41.53.

(21) Takahashi, N.; Suzuki, A.; Kimura, Y.; Miyamoto, S.; Tamura, S. Structure of piericidin B and stereochemistry of piericidins. Tetrahedron Lett. 1967, 8 (21), 1961–1964. DOI: 10.1016/s0040-4039(00)90764-0.

(22) Matsumoto, M.; Mogi, K.; Nagaoka, K.; Ishizeki, S.; Kawahara, R.; Nakashima, T. New piericidin glucosides, glucopiericidins A and B. J. Antibiot. 1987, 40 (2), 149–156. DOI: 10.7164/antibiotics.40.149.

(23) Shaaban, K. A.; Helmke, E.; Kelter, G.; Fiebig, H. H.; Laatsch, H. Glucopiericidin C: a cytotoxic piericidin glucoside antibiotic produced by a marine Streptomyces isolate. J. Antibiot. 2011, 64 (2), 205. DOI: 10.1038/ja.2010.125.

(24) Hall, C.; Wu, M.; Crane, F. L.; Takahashi, H.; Tamura, S.; Folkers, K. Piericidin A: a new inhibitor of mitochondrial electron transport. Biochem. Biophys. Res. Commun. 1966, 25 (4), 373–377. DOI: 10.1016/0006-291x(66)90214-2.

(25) Darrouzet, E.; Issartel, J.-P.; Lunardi, J.; Dupuis, A. The 49-kDa subunit of NADH-ubiquinone oxidoreductase (Complex I) is involved in the binding of piericidin and rotenone, two quinone-related inhibitors. FEBS Lett. 1998, 431 (1), 34–38. DOI: 10.1016/S0014-5793(98)00719-4.

(26) Hwang, J. H.; Kim, J. Y.; Cha, M. R.; Ryoo, I. J.; Choo, S. J.; Cho, S. M.; Tsukumo, Y.; Tomida, A.; Shin-Ya, K.; Hwang, Y. I.; Yoo, I. D.; Park, H. R. Etoposide-resistant HT-29 human colon carcinoma cells during glucose deprivation are sensitive to piericidin A, a GRP78 down-regulator. J. Cell Physiol. 2008, 215 (1), 243–250. DOI: 10.1002/jcp.21308.

(27) Horgan, D. J.; Ohno, H.; Singer, T. P.; Casida, J. E. Studies on the respiratory chain-linked reduced nicotinamide adenine dinucleotide dehydrogenase. XV. Interactions of piericidin with the mitochondrial respiratory chain. J. Biol. Chem. 1968, 243 (22), 5967–5976. DOI: 10.1016/S0021-9258(18)94515-1.

(28) Yoshida, S.; Nagao, Y.; Watanabe, A.; Takahashi, N. Structure-activity relationship in piericidins inhibitors on the electron transport system in mitochondria. Agric. Biol. Chem. 1980, 44 (12), 2921–2924. DOI: 10.1080/00021369.1980.10864438.

(29) Chung, K. H.; Cho, K. Y.; Asami, Y.; Takahashi, N.; Yoshida, S. New 4-hydroxypyridine and 4-hydroxyquinoline derivatives as inhibitors of NADH-ubiquinone reductase in the respiratory chain. ZNC 1989, 44 (7-8), 609–616. DOI: 10.1515/znc-1989-7-811.

(30) Liu, Q.; Yao, F.; Chooi, Y. H.; Kang, Q.; Xu, W.; Li, Y.; Shao, Y.; Shi, Y.; Deng, Z.; Tang, Y.; You, D. Elucidation of Piericidin A1 biosynthetic locus revealed a thioesterase-dependent mechanism of alpha-pyridone ring formation. Chem. Biol. 2012, 19 (2), 243–253. DOI: 10.1016/j.chembiol.2011.12.018.

(31) Chen, Y.; Zhang, W.; Zhu, Y.; Zhang, Q.; Tian, X.; Zhang, S.; Zhang, C. Elucidating hydroxylation and methylation steps tailoring piericidin A1 biosynthesis. Org. Lett. 2014, 16 (3), 736–739. DOI: 10.1021/ol4034176.

(32) Li, Y.; Kong, L.; Shen, J.; Wang, Q.; Liu, Q.; Yang, W.; Deng, Z.; You, D. Characterization of the positive SARP family regulator PieR for improving piericidin A1 production in Streptomyces piomogeues var. Hangzhouwanensis. Synth. Syst. Biotechnol. 2019, 4 (1), 16–24. DOI: 10.1016/j.synbio.2018.12.002.

(33) https://mira-assembler.sourceforge.net/docs/DefinitiveGuideToMIRA.html.

(34) Prjibelski, A.; Antipov, D.; Meleshko, D.; Lapidus, A.; Korobeynikov, A. Using SPAdes De Novo Assembler. Curr. Protoc. Bioinform. 2020, 70 (1), e102. DOI: 10.1002/cpbi.102.

(35) Wick, R. R.; Judd, L. M.; Gorrie, C. L.; Holt, K. E. Unicycler: Resolving bacterial genome assemblies from short and long sequencing reads. PLoS Comput. Biol. 2017, 13 (6), e1005595. DOI: 10.1371/journal.pcbi.1005595.

(36) Kolmogorov, M.; Bickhart, D. M.; Behsaz, B.; Gurevich, A.; Rayko, M.; Shin, S. B.; Kuhn, K.; Yuan, J.; Polevikov, E.; Smith, T. P. L.; Pevzner, P. A. metaFlye: scalable long-read metagenome assembly using repeat graphs. Nat. Methods 2020, 17 (11), 1103–1110. DOI: 10.1038/s41592-020-00971-x.

(37) Walker, B. J.; Abeel, T.; Shea, T.; Priest, M.; Abouelliel, A.; Sakthikumar, S.; Cuomo, C. A.; Zeng, Q.; Wortman, J.; Young, S. K.; Earl, A. M. Pilon: an integrated tool for comprehensive microbial variant detection and genome assembly improvement. PLoS One 2014, 9 (11), e112963. DOI: 10.1371/journal.pone.0112963.

(38) Blin, K.; Shaw, S.; Augustijn, H. E.; Reitz, Z. L.; Biermann, F.; Alanjary, M.; Fetter, A.; Terlouw, B. R.; Metcalf, W. W.; Helfrich, E. J. N.; van Wezel, G. P.; Medema, M. H.; Weber, T. antiSMASH 7.0: new and improved predictions for detection, regulation, chemical structures and visualisation. Nucleic Acids Res. 2023, 51 (W1), W46–W50. DOI: 10.1093/nar/gkad344.

(39) Li, W.; O’Neill, K. R.; Haft, D. H.; DiCuccio, M.; Chetvernin, V.; Badretdin, A.; Coulouris, G.; Chitsaz, F.; Derbyshire, M. K.; Durkin, A. S.; Gonzales, N. R.; Gwadz, M.; Lanczycki, C. J.; Song, J. S.; Thanki, N.; Wang, J.; Yamashita, R. A.; Yang, M.; Zheng, C.; Marchler-Bauer, A.; Thibaud-Nissen, F. RefSeq: expanding the Prokaryotic Genome Annotation Pipeline reach with protein family model curation. Nucleic Acids Res 2021, 49 (D1), D1020–D1028. DOI: 10.1093/nar/gkaa1105.

(40) Dashti, Y.; Nakou, I. T.; Mullins, A. J.; Webster, G.; Jian, X.; Mahenthiralingam, E.; Challis, G. L. Discovery and biosynthesis of bolagladins: unusual lipodepsipeptides from Burkholderia gladioli clinical isolates. Angew. Chem. Int. Ed. 2020, 59 (48), 21553–21561. DOI: 10.1002/anie.202009110.

(41) Wilkinson, C. J.; Hughes-Thomas, Z. A.; Martin, C. J.; Böhm, I.; Mironenko, T.; Deacon, M.; Wheatcroft, M.; Wirtz, G.; Staunton, J.; Leadlay, P. F. Increasing the efficiency of heterologous promoters in actinomycetes. J. Mol. Microbiol. Biotechnol. 2002, 4 (4), 417–426.

(42) Sumang, F. A.; Errington, J.; Dashti, Y. Biosynthesis of the quinovosamycin nucleoside antibiotics diverges from that of tunicamycins by additional sugar processing genes. Bioorg. Chem. 2025, 160, 108431. DOI: 10.1016/j.bioorg.2025.108431.

(43) Amalin, D. M.; Vasquez, E. A. A handbook on Philippine sweet potato arthropod pests and their natural enemies; International Potato Center, 1993.

(44) Carballo, J. L.; Hernández-Inda, Z. L.; Pérez, P.; García-Grávalos, M. D. A comparison between two brine shrimp assays to detect in vitrocytotoxicity in marine natural products. BMC Biotechnol. 2002, 2 (1), 17. DOI: 10.1186/1472-6750-2-17.

(45) Sakuma, M. Probit analysis of preference data. Appl. Entomol. Zool. 1998, 33 (3), 339–347. DOI: 10.1303/aez.33.339.

