## Supplementary Information for "An *O*-acetylated derivative of piericidin A1 produced by *Kitasatospora* sp. A2-31 has potent activity against the cacao mirid bug, *Helopeltis bakeri* Poppius"

[1] Department of Chemistry, University of Warwick, Coventry CV4 7AL, United Kingdom
[2] School of Medical Sciences, Faculty of Medicine and Health, University of Sydney, Sydney NSW 2015, Australia
[3] Sydney Infectious Diseases Institute, University of Sydney, Sydney NSW 2015, Australia
[4] National Institute of Molecular Biology and Biotechnology (BIOTECH), University of the Philippines Los Baños, College, Laguna, Philippines
[5] Warwick Integrative Synthetic Biology Centre, University of Warwick, Coventry CV4 7AL, United Kingdom
[6] School of Biology, Newcastle University, Newcastle upon Tyne, UK
[7] Department of Biology, College of Science, De La Salle University, Manila, Philippines
[8] Center for Natural Science and Environmental Research, De La Salle University, Manila Philippines
[9] Department of Biochemistry and Molecular Biology, Monash University, Clayton, Victoria 3800, Australia
[10] ARC Centre of Excellence for Innovations in Peptide and Protein Science, Monash University, Clayton, Victoria 3800, Australia

[‡]Present Address: School of Medical Sciences, Faculty of Medicine and Health, University of Sydney, Sydney NSW 2015, Australia (F.A.S.); UCB Pharma Ltd, 208 Bath Road, Slough, Berkshire, SL1 3WE, United Kingdom (E.L.C.d.l.S.).


| **Contents** | **Page** |
|---|---|
| Figure S1. Cacao mirid bug and their damage symptoms on cacao pod. | S2 |
| Figure S2. Structure of acetylpiericidin A1 with numbering used in NMR table S1. | S3 |
| Table S1. $^{1}$H and $^{13}$C NMR chemical shifts of acetylpiericidin A1 in DMSO-$d_6$. | S3 |
| Figure S3. $^{1}$H NMR spectrum of acetylpiericidin A1 in DMSO-$d_6$. | S4 |
| Figure S4. $^{13}$C NMR spectrum of acetylpiericidin A1 in DMSO-$d_6$. | S5 |
| Figure S5. COSY spectrum of acetylpiericidin A1 in DMSO-$d_6$. | S6 |
| Figure S6. HSQC spectrum of acetylpiericidin A1 in DMSO-$d_6$. | S7 |
| Figure S7. HMBC spectrum of acetylpiericidin A1 in DMSO-$d_6$. | S8 |
| Table S2. Genes in the acetylpiericidin A1 biosynthetic gene cluster and percentage identity of the proteins they encode to proteins of known function. | S9 |
| Figure S8. Multiple sequence alignment of ApiF with spermidine *N*-acetyltransferase SpeG and mass spectrum of expressed ApiF | S10 |

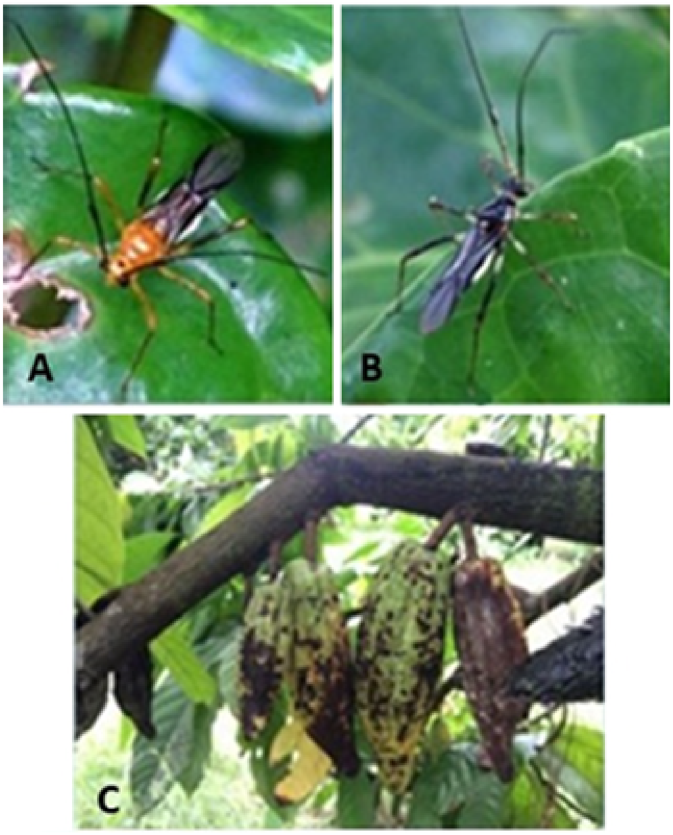


**Figure S1.** Cacao mirid bug, A) adult female, B) adult male, and C) damage symptoms on cacao pod

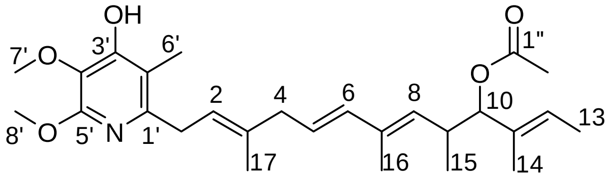


**Figure S2.** Structure of acetylpiericidin A1 with numbering used in NMR table S1.

**Table S1.** $^{1}$H and $^{13}$C NMR chemical shifts of acetylpiericidin A1 in DMSO-$d_6$.

| Position | $\delta_C$ | $\delta_H$ | HMBC |
|---|---|---|---|
| 1 | 33.7 | 3.27, d ($J$ = 6.9) | C2, C3, C2', C3' |
| 2 | 122.2 | 5.30, dd ($J$ = 15.1, 7.8 ) | C1, C4, C17, C2' |
| 3 | 133.7 | | |
| 4 | 42.2 | 2.72, d ($J$ = 7.0) | C2, C6, C17 |
| 5 | 125.6 | 5.53, m | C3, C4, C7 |
| 6 | 135.7 | 5.99, d ($J$ = 15.5) | C4, C8, C16 |
| 7 | 133.2 | | |
| 8 | 132.6 | 5.11, d ($J$ = 9.5) | C6, C9, C15, C16 |
| 9 | 34.3 | 2.80, m | C7, C10, C15 |
| 10 | 82.4 | 4.87, d ($J$ = 8.5) | C8, C9, C11, C12, C14, C15, C1" |
| 11 | 132.5 | | |
| 12 | 123.6 | 5.45, m | C10, C11, C13, C14 |
| 13 | 11.8 | 1.56, d ($J$ = 8.8) | C11, C12 |
| 14 | 12.0 | 1.57, s | C10, C11, C12 |
| 15 | 16.9 | 0.80, d ($J$ = 6.8) | C8, C9, C10 |
| 16 | 12.3 | 1.69, s | C6, C7, C8 |
| 17 | 16.1 | 1.69, s | C2, C3, C4 |
| 1′ | 150.0 | | |
| 2′ | 112.7 | | |
| 3′ | 155.2 | -OH, 9.78, s | C2', C3', C4' |
| 4′ | 128.5 | | |
| 5′ | 154.5 | | |
| 6′ | 10.3 | 1.97, s | C1', C2', C3' |
| 7′ | 60.0 | 3.63, s | C4' |
| 8′ | 52.5 | 3.80, s | C5' |
| 1″ | 169.5 | | |
| 2″ | 20.4 | 1.87, s | C1" |

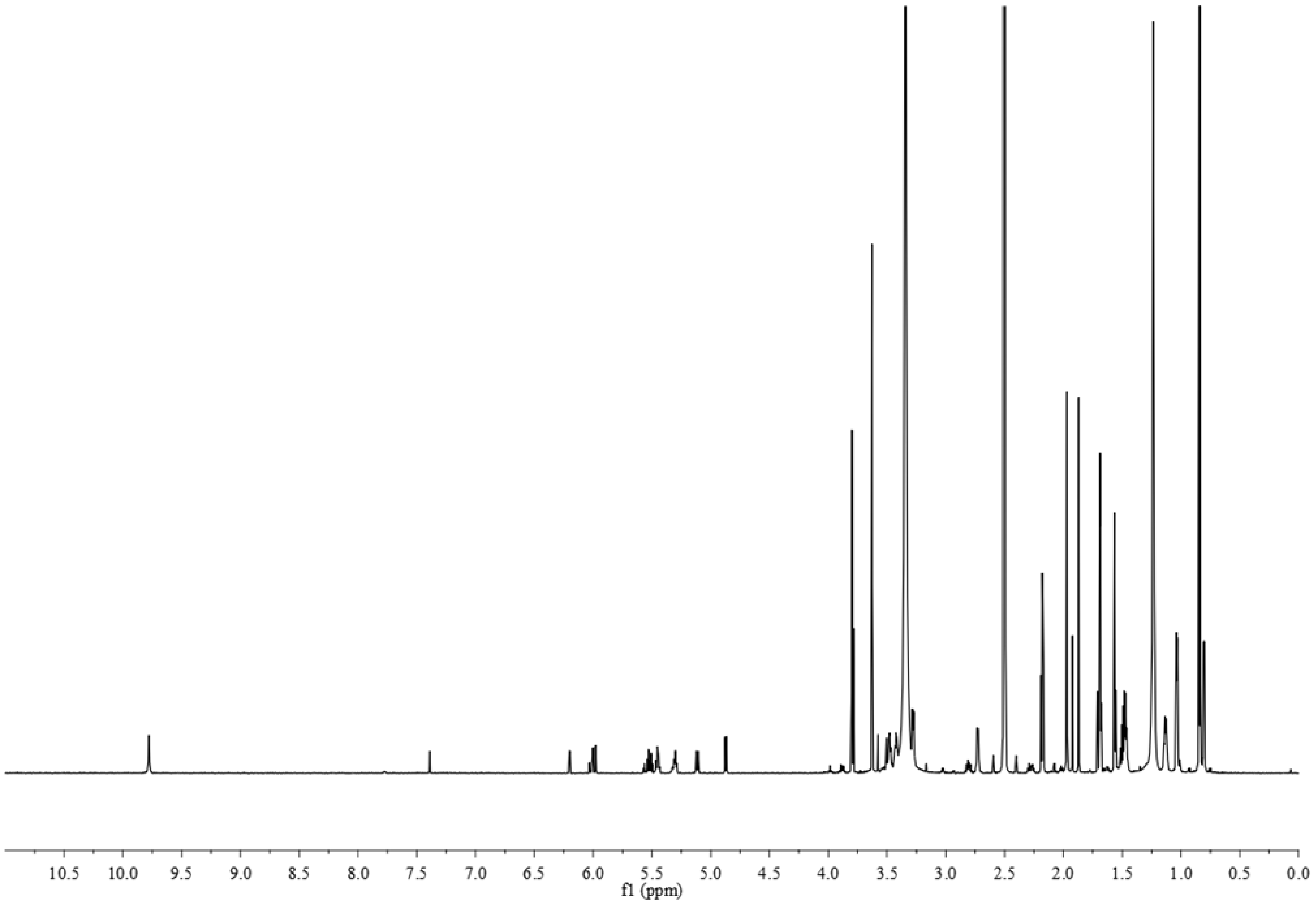


**Figure S3.** $^1$H NMR spectrum of acetylpiericidin A1 in DMSO-$d_6$.

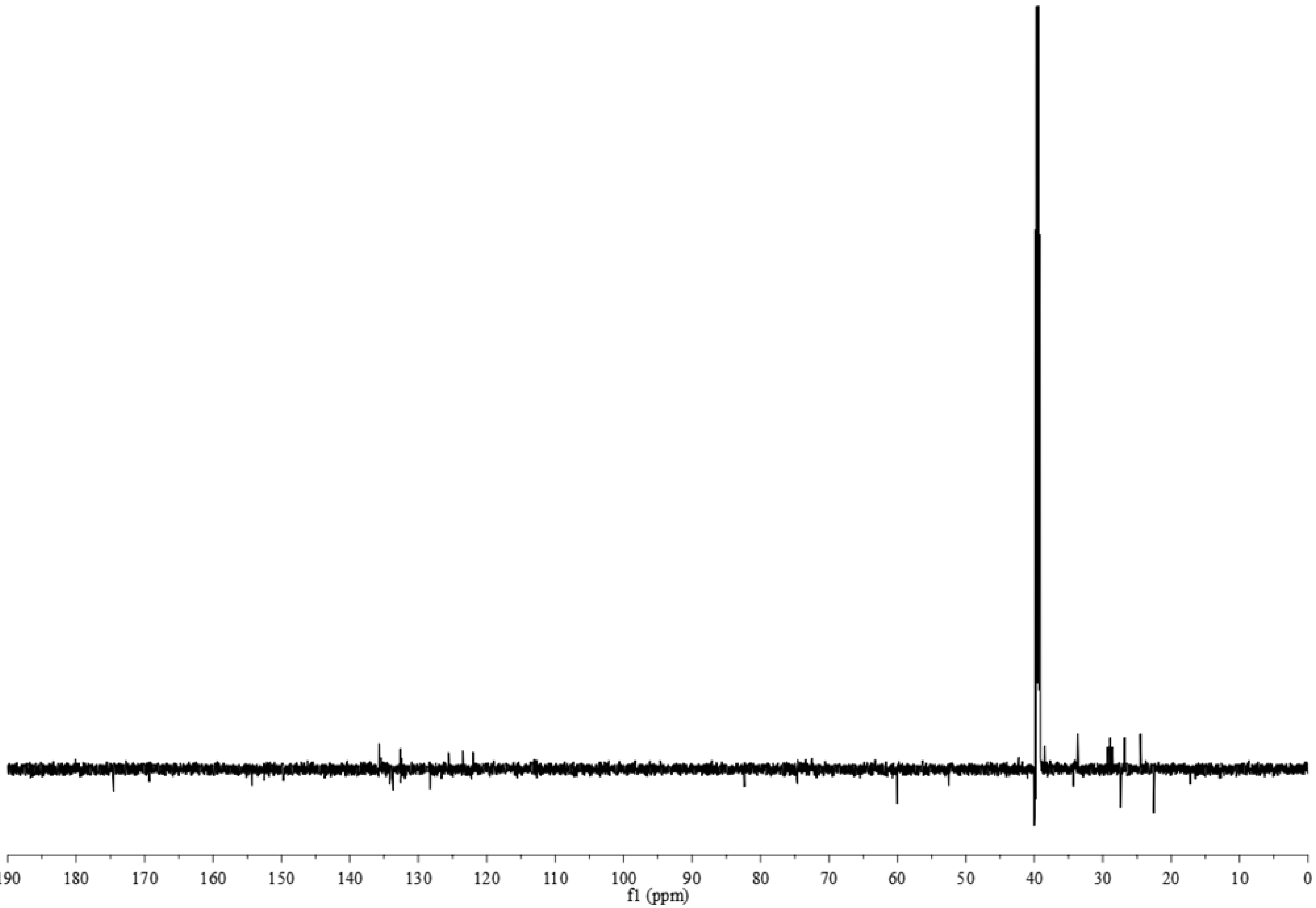


**Figure S4.** $^{13}$C NMR spectrum of acetylpiericidin A1 in DMSO-$d_6$.

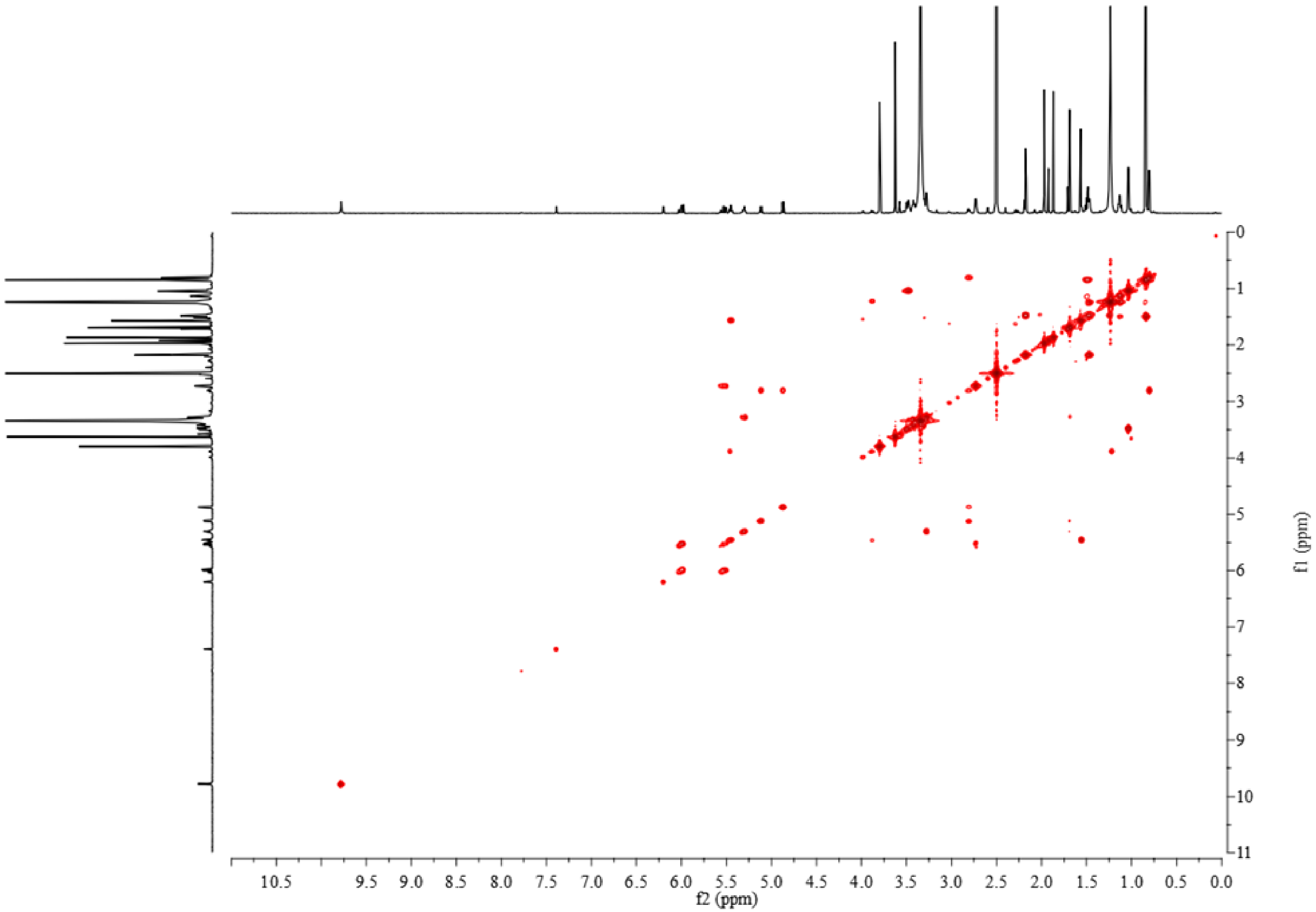


**Figure S5.** COSY spectrum of acetylpiericidin A1 in DMSO-$d_6$.

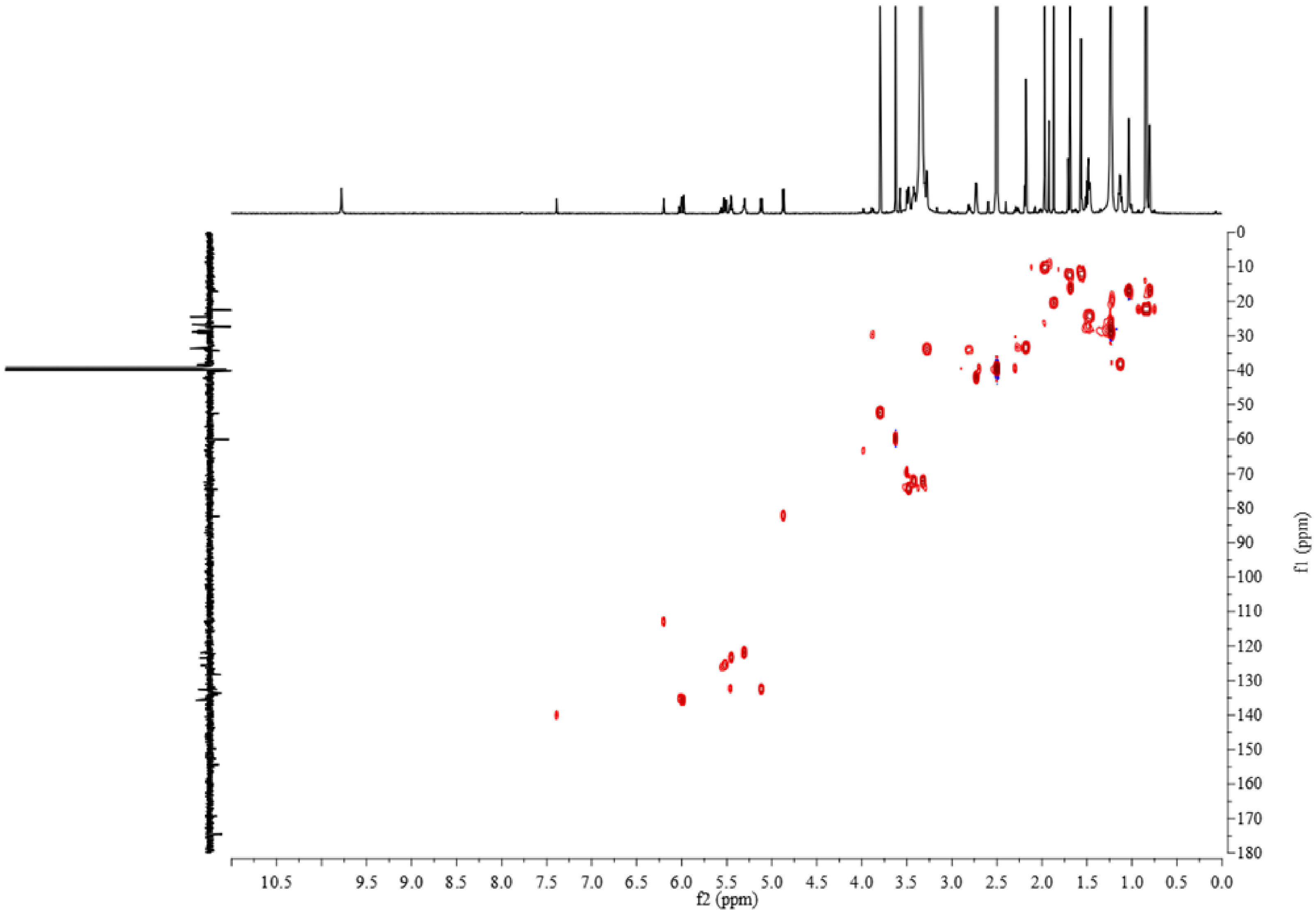


**Figure S6.** HSQC spectrum of acetylpiericidin A1 in DMSO-$d_6$.

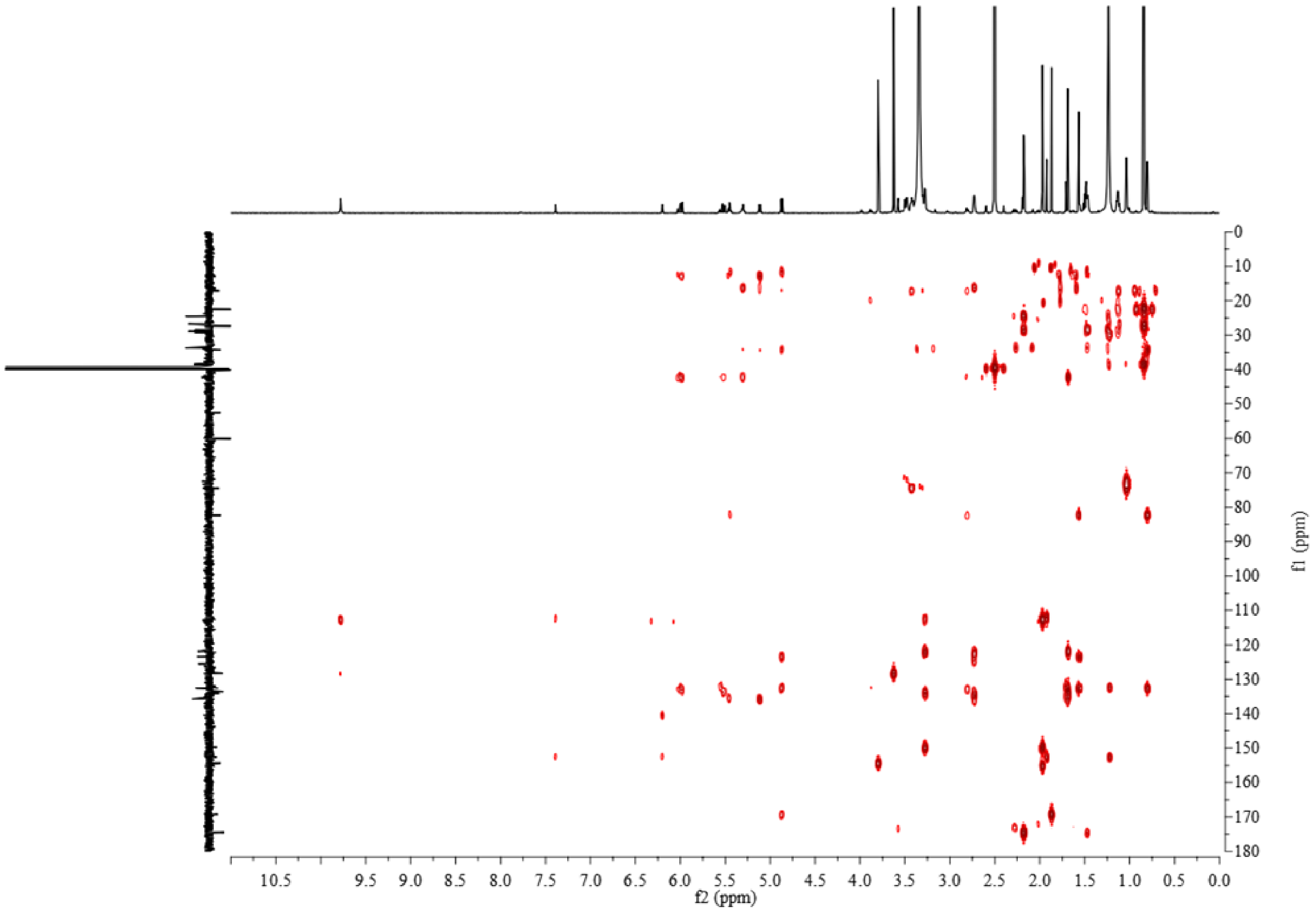


**Figure S7.** HMBC spectrum of acetylpiericidin A1 in DMSO-$d_6$.

**Table S2.** Genes in the acetylpiericidin A1 biosynthetic gene cluster and percentage identity of the proteins they encode to proteins of known function.

| Gene/Protein | Length bp/aa | Similar Proteins | % aa Identity |
|---|---|---|---|
| *apiE*/ApiE | 1434/478 | PieE (*Streptomyces piomogenus*) | 97 |
| *apiB2*/ApiB2 | 774/257 | PieB2 (*Streptomyces piomogenus*) | 99 |
| *apiD*/ApiD | 1842/ 613 | PieD (*Streptomyces piomogenus*) | 98 |
| *apiC*/ApiC | 513/170 | PieC (*Streptomyces piomogenus*) | 98 |
| *apiB1*/ApiB1 | 687/ 228 | PieB1 (*Streptomyces piomogenus*) | 99 |
| *apiA6*/ApiA6 | 7071/2357 | PieA6 (*Streptomyces piomogenus*) | 91 |
| *apiA5*/ApiA5 | 5592/1864 | PieA5 (*Streptomyces piomogenus*) | 93 |
| *apiA4*/ApiA4 | 6486/2161 | PieA4 (*Streptomyces piomogenus*) | 92 |
| *apiA3*/ApiA3 | 5151/1717 | PieA3 (*Streptomyces piomogenus*) | 92 |
| *apiA2*/ApiA2 | 10149/3383 | PieA2 (*Streptomyces piomogenus*) | 93 |
| *apiA1*/ApiA1 | 7773/2791 | PieA1 (*Streptomyces piomogenus*) | 88 |
| *apiR*/ApiR | 723/241 | PieR (*Streptomyces piomogenus*) | 85 |
| *apiF*/ApiF | 531/176 | GNAT family acetyltransferase (*Streptomyces novaecaesareae*) | 83 |

**a**

```
SpeG      --MNSQLTLRALERGDLRFIHN------LNNNRNIMSYWFEEPYESFDELEELYNKHIHD      52
ApiF      MLQGAHITLRARHETDVPVLQAELYDDVATRSRADSRAWRPIPPGSTQSP--YAVSGASD      58
             .:::****  ..  *:  .::          ...*      *    *   *  :.        .    *

SpeG      NAERRFVVEDAQKNLIGLVELIEINYIHRSAEFQIIIAPEHQGKGFARTLINRALDYSFT      112
ApiF      EAACFSVVETASGELAGEALLWGIDTHNRTAHLGIALRPAHRGRGFAADVLRVLCRFGFT      118
          :*     *** *.  :* *  .  *   *:   :*:*.:  *  :  *  *:*:***   ::.      :.**

SpeG      ILNLHKIYLHVAVENPKAVHLYEECGFVEEGHLVEEFFINGRYQDVKRMYILQSKYLNRS      172
ApiF      VLGLNRLQLETLADNAPMVRAATAAGFTTEGTLRRSAWACGEFADQVVLGLLAEEWRG--      176
          :*.*::: *..  .:*    *:      .**.  ** *  ..  :   *.:  *     :  :*  .::  .

SpeG      E      173
ApiF      -      176
```

**b**

MGSSHHHHHHSSGLVPRGSHMMLQGAHITLRARHETDVPVLQAELYDDVATRSRA
DSRAWRPIPPGSTQSPYAVSGASDEAACFSVVETASGELAGEALLWGIDTHNRTAHL
GIALRPAHRGRGFAADVLRVLCRFGFTVLGLNRLQLETLADNAPMVRAATAAGFTT
EGTLRRSAWACGEFADQVVLGLLAEEWRG

**c**

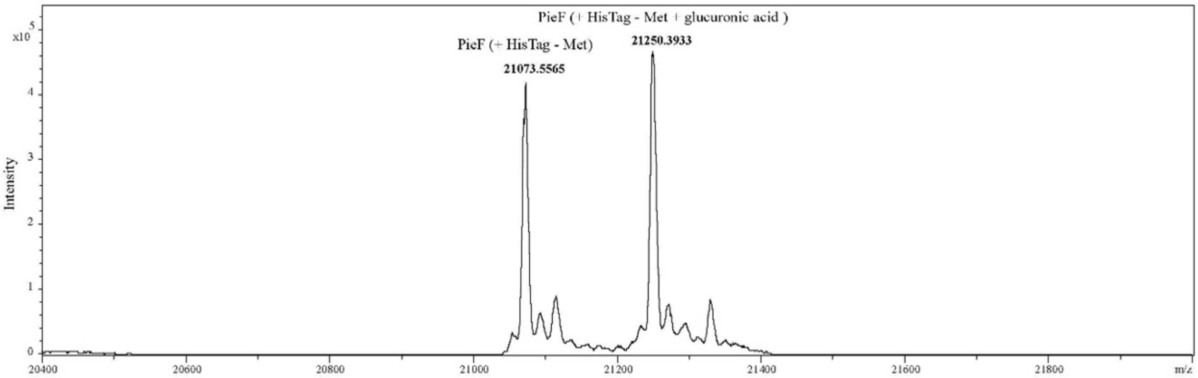


Expected Mass: ApiF (+ HisTag): 21204 Da
Observed Mass 1: ApiF (+ HisTag - Met ): 21073 Da
Observed Mass 2: ApiF (+ HisTag - Met + glucuronic acid): 21250 Da

**Figure S8. a)** Multiple sequence alignment of ApiF with spermidine *N*-acetyltransferase SpeG. **b)** Amino acid sequence of expressed ApiF linked to HisTag. **c)** Mass spectrum of expressed product shows that methionine is cleaved from protein and also shows glucuronidation of ApiF.
